# Monitoring the 5-methoxycarbonylmethyl-2-thiouridine (mcm5s2U) modification in eukaryotic tRNAs via the gamma-toxin endonuclease

**DOI:** 10.1101/253708

**Authors:** Jenna M. Lentini, Jillian Ramos, Dragony Fu

## Abstract

The post-transcriptional modification of tRNA at the wobble position is a universal process occurring in all domains of life. In eukaryotes, the wobble uridine of particular tRNAs is transformed to the 5-methoxycarbonylmethyl-2-thiouridine (mcm5s2U) modification which is critical for proper mRNA decoding and protein translation. However, current methods to detect mcm5s2U are technically challenging and/or require specialized instrumental expertise. Here, we show that gamma-toxin endonuclease from the yeast *Kluyveromyces lactis* can be used as a probe for assaying mcm5s2U status in the tRNA of diverse eukaryotic organisms ranging from protozoans to mammalian cells. The assay couples the mcm5s2U-dependent cleavage of tRNA by gamma-toxin with standard molecular biology techniques such as Northern blot analysis or quantitative PCR to monitor mcm5s2U levels in multiple tRNA isoacceptors. The results gained from the gamma-toxin assay reveals the evolutionary conservation of the mcm5s2U modification across eukaryotic species. Moreover, we have employed the gamma-toxin assay to verify uncharacterized eukaryotic Trm9 and Trm112 homologs that catalyze the formation of mcm5s2U. These findings demonstrate the use of gamma-toxin as a detection method to monitor mcm5s2U status in diverse eukaryotic cell types for cellular, genetic and biochemical studies.

## INTRODUCTION

The post-transcriptional modification of uridine at the wobble position of tRNA is ubiquitous in all organisms and plays a critical role in proper mRNA decoding during translation (reviewed in (Grosjean et al. 2010; Agris et al. 2017; Schaffrath and Leidel 2017)). In eukaryotes, the wobble uridine of certain tRNAs can undergo a series of modifications to generate the modified nucleotide, methoxycarbonylmethyl-2-thiouridine (mcm5s2U) (Figure 1A). While the specific order of biochemical intermediates leading to mcm5s2U remains to be clarified (Songe-Moller et al. 2010; Chen et al. 2011a; van den Born et al. 2011), several proteins have been identified in the yeast *S. cerevisiae* that are required for mcm5s2U formation in tRNA. The multi-subunit Elongator complex is required for early steps of mcm5U formation through the generation of 5-carbonylmethyluridine (cm5U) and 5-carbamoylmethyluridine (ncm5U) via a proposed enzymatic reaction involving acetyl-CoA (reviewed in (Karlsborn et al. 2014b; Dauden et al. 2017; Kolaj-Robin and Seraphin 2017)). The cm5U and/or ncm5U modification is then further modified by a heterodimeric methyltransferase complex consisting of the Trm9 enzymatic subunit and the Trm112 structural protein (Kalhor and Clarke 2003; Mazauric et al. 2010; Liger et al. 2011; Letoquart et al. 2015; Bourgeois et al. 2017). In a subset of tRNAs, the mcm5U base can undergo even further modification by the Nnc2-Ncs6 thiolase to generate the final mcm5s2U modification (Dewez et al. 2008; Huang et al. 2008; Leidel et al. 2009; Noma et al. 2009; Nakai et al. 2017).

**Figure 1.**
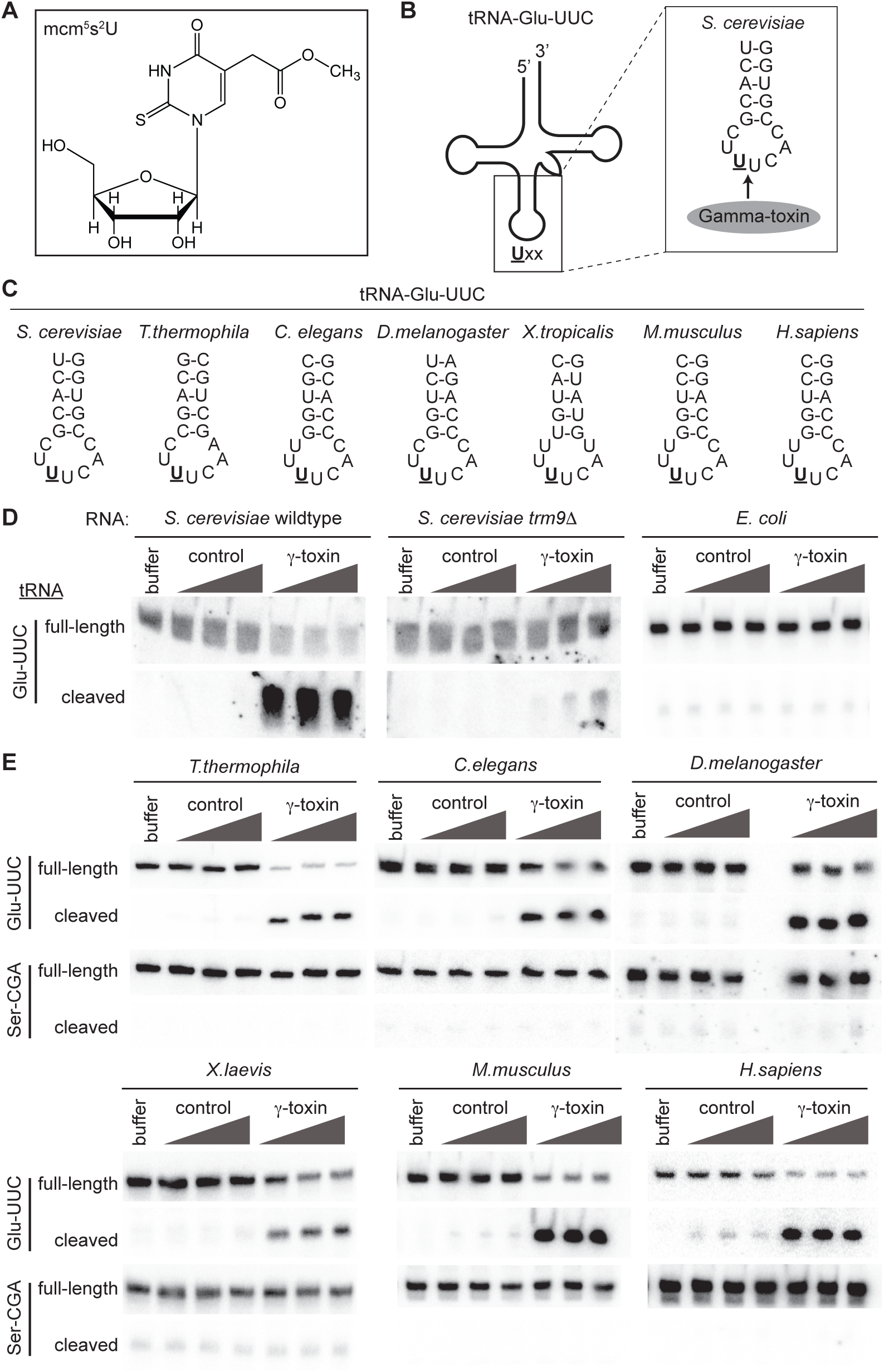
Purified gamma-toxin can cleave mcm5s2U containing tRNA species from a wide range of organisms. (A) Chemical structure of the mcm5s2U modification found on the wobble Uridine of select tRNA species. (B) Schematic of *S. cerevisiae* tRNA-Glu-UUC highlighting the anticodon stem and position of Gamma-toxin cleavage 3’ of the mcm5s2U modification denoted by <u>U</u>. (C) Anticodon stems of tRNA-Glu-UUC from Eukaryotic organisms tested in this study. The underlined <u>U</u> denotes the mcm5s2U modification. (D) Northern blots of *S. cerevisiae* or *E.coli* total RNA pre-incubated with buffer or increasing amounts of either a control purification eluate or gamma-toxin. The blot was hybridized with a probe against tRNA-Glu-UUC to detect full-length or cleaved tRNA products (E) Total RNA from the indicated Eukaryotic organisms was incubated with either buffer, control purification eluate or purified gamma-toxin followed by Northern blot analysis as in (D). The blot was probed for tRNA-Ser-CGA as a specificity and loading control.

The mcm5s2U modification plays a key role in the efficiency and fidelity of mRNA translation to modulate protein levels in eukaryotes (Huang et al. 2005; Johansson et al. 2008; Bauer et al. 2012; Patil et al. 2012a; Fernandez-Vazquez et al. 2013; Rezgui et al. 2013; Deng et al. 2015; Klassen et al. 2015; Tukenmez et al. 2015; Klassen et al. 2016a). Due to its effects on protein synthesis, the presence of mcm5s2U at the wobble position is crucial for proper cellular growth, metabolism and organismal development (Esberg et al. 2006; Bjork et al. 2007; Lin et al. 2013; Leitner et al. 2015; Karlsborn et al. 2016; Klassen et al. 2016b; Schaffrath and Klassen 2017). Moreover, the mcm5U-class of tRNA modifications has been shown to play a critical role in the cellular stress response by regulating the translation of proteins involved in redox homeostasis, DNA repair and telomere silencing (Begley et al. 2007; Chan et al. 2010; Fu et al. 2010a; Chen et al. 2011b; Patil et al. 2012b; Begley et al. 2013; Fernandez-Vazquez et al. 2013; Zinshteyn and Gilbert 2013; Endres et al. 2015). Significantly, the mcm5s2U modification is also required for proper neurodevelopment in worms, mice and humans since gene mutations that reduce mcm5s2U levels lead to a spectrum of neurological disorders including familial dysautonomia (Chen et al. 2009; Karlsborn et al. 2014a; Yoshida et al. 2015; Kojic and Wainwright 2016). Thus, the detection of mcm5s2U levels will be critical for understanding the role of tRNA modifications in diverse biological processes.

The use of thin layer chromatography (TLC) for separation of digested RNA nucleosides has been a classic technique for identifying and detecting modifications such as mcm5s2U (Grosjean et al. 2004). While TLC analysis remains a viable approach for analysis of mcm5s2U, the radiolabeling, digestion, spotting and chromatographic separation of RNA on TLC plates is labor-intensive and time-consuming. In addition, the resolution of mcm5s2U by 2-dimensional TLC requires extensive optimization for clear interpretation (Grosjean et al. 2007).

High pressure-liquid chromatography (HPLC) provides an alternative method for detection of mcm5s2U through the separation of nucleosides generated by RNA digestion (Huang et al. 2005; Mazauric et al. 2010; Chen et al. 2011a). Furthermore, HPLC has been coupled with mass spectrometry (HPLC-MS) for sensitive identification and quantification of RNA modifications (Su et al. 2014; Cai et al. 2015; Cao and Limbach 2015; Basanta-Sanchez et al. 2016; Ross et al. 2016). However, HPLC-MS requires lengthy run times, extensive optimization and the generation of mcm5s2U standards. Thus, the expertise and instrumentation costs associated with HPLC-MS instrumentation is beyond the capabilities of many laboratories wishing to study tRNA modifications such as mcm5s2U.

Many tRNA modifications can also be detected by reverse transcriptase (RT)-based primer extension methods, differential probe hybridization or reactivity to certain chemical reagents (reviewed in (Schwartz and Motorin 2016; Helm and Motorin 2017)). However, the mcm5s2U modification has no detectable effect on oligonucleotide hybridization, RT extension or nucleotide misincorporation (Hiley et al. 2005; Motorin et al. 2007; Behm-Ansmant et al. 2011). Loss of the thiol group in mcm5s2U can be detected by differential tRNA migration in the presence of acryloylaminophenylmercuric chloride but changes in the methylester group are undetectable using the same technique (Igloi 1992; Kaneko et al. 2003). While chemical treatment of tRNA with low concentrations of NaOH can remove the methyl group from mcm5s2U to generate cm5U, this treatment is complete, leads to RNA degradation, and does not lead to any change in RNA migration or RT primer extension (Mazauric et al. 2010; Chen et al. 2011a; Leihne et al. 2011). Moreover, there are no known published antibodies that have been generated against mcm5s2U.

Due to the reasons noted above, we sought to develop an assay for detection of mcm5s2U that would utilize standard molecular biology assays and obviate the need for TLC or HPLC-MS. We focused our attention on gamma-toxin, a ribonuclease secreted from the milk yeast *Kluyveromyces lactis* that catalyzes the endonucleolytic cleavage of foreign yeast tRNAs in a mcm5s2U-dependent manner (reviewed in (Lu et al. 2005; Jablonowski et al. 2006; Jablonowski and Schaffrath 2007; Lu et al. 2008)). Gamma-toxin cleaves tRNAs-Glu-UUC, Lys-UUU and Gln-UUG between positions 34 and 35 of the anticodon loop with cleavage being highly dependent upon the presence of the mcm5s2U modification (Figure 1B, tRNA-Glu-UUC shown). *S.cerevisiae* yeast cells that are deficient for any of the enzymes that form mcm5s2U in tRNA are resistant to gamma toxin-mediated tRNA cleavage and cell death (Frohloff et al. 2001; Fichtner and Schaffrath 2002; Jablonowski et al. 2006; Huang et al. 2008; Studte et al. 2008). Gamma-toxin has been used in *S. cerevisiae* to uncover proteins and pathways involved in mcm5s2U modification (Huang et al. 2008; Studte et al. 2008; Mehlgarten et al. 2009; Mehlgarten et al. 2010; Judes et al. 2016; Mehlgarten et al. 2017). Based upon the sequence requirements and substrate specificity of gamma-toxin, we predicted that gamma-toxin could be used to cleave mcm5s2U-containing tRNAs isolated from other eukaryotic organisms. Here, we show that gamma-toxin can be employed as a molecular tool for monitoring mcm5s2U status across a variety of eukaryotic species, including model eukaryotic organisms. Moreover, we demonstrate that gamma-toxin can be used for the discovery and *in vitro* validation of uncharacterized eukaryotic proteins involved in the formation of mcm5s2U.

## RESULTS and DISCUSSION

### Gamma-toxin cleaves mcm5s2U-containing tRNAs from diverse eukaryotic organisms

Due to the specificity of gamma-toxin for yeast tRNAs containing mcm5s2U, we hypothesized that gamma-toxin could be used to monitor mcm5s2U status in additional eukaryotic organisms besides *S. cerevisiae* (Figure 1C). To assay for tRNA cleavage, we incubated recombinant gamma-toxin purified from *E. coli* with total RNA isolated from different eukaryotic organisms followed by Northern blot hybridization analysis. To monitor non-specific RNA cleavage due to possible contaminating nucleases, we also incubated RNA with control purifications from *E. coli* cells containing vector alone. As a positive control, we tested the activity of gamma-toxin against *S. cerevisiae* tRNA-Glu-UUC, the primary physiological target of gamma-toxin endonuclease (Lu et al. 2005; Jablonowski et al. 2006). No tRNA cleavage products were detected when *S. cerevisiae* total RNA was incubated with buffer or increasing amounts of a control protein purification from *E. coli* cells (Figure 1D, *S. cerevisiae* wildtype, buffer and control lanes). However, incubation of *S. cerevisiae* RNA with purified gamma-toxin led to the cleavage of tRNA-Glu-UUC and the accumulation of cleaved tRNA fragments (Figure 1D, *S. cerevisiae* wildtype + γ-toxin). Gamma-toxin cleavage of *S. cerevisiae* tRNA-Glu-UUC was greatly reduced with RNA extracted from *trm9*Δ cells, which are unable to form the mcm5s2U modification (Figure 1D, *trm9*Δ). Moreover, gamma-toxin was unable to cleave *E. coli* tRNA-Glu-UUC, which contains a 5-methylaminomethyluridine (mnm5U) modification instead of mcm5s2U (Figure 1D, *E. coli* + γ-toxin).

Upon confirming the activity of gamma-toxin, we tested whether gamma-toxin displays endonuclease activity against mcm5s2U-containing tRNAs of additional eukaryotic organisms. Remarkably, we find that gamma-toxin can efficiently and specifically cleave tRNA-Glu-UUC isolated from a diversity of eukaryotes including *Tetrahymena thermophila* (ciliated protozoa), *Caenorhabditis elegans* (nematode), *Drosophila melanogaster* (fruit fly), *Xenopus tropicalis* (frog), *Mus musculus* (mouse embryonic fibroblasts) and *Homo sapiens* (human embryonic fibroblast cells) (Figure 1E, Glu-UUC + γ-toxin). In addition to the organisms described here, we have also used gamma-toxin to demonstrate the presence of mcm5s2U in tRNA-Glu-UUC of the parasitic protozoan, *Toxoplasma gondii* (Padgett et al. 2018). In all cases where total eukaryotic RNA was incubated with gamma-toxin, we detected tRNA fragments indicative of only a single tRNA cleavage event within the anticodon loop of tRNA-Glu-UUC without any other products of non-specific degradation (Figure 1E, Glu-UUC, data not shown). In contrast to tRNA-Glu-UUC of each eukaryote, we detected no detectable cleavage of tRNA-Ser-CGA, which does not contain a wobble uridine and hence, lacks the mcm5s2U modification (Figure 1E, Ser-CGA). Our results demonstrate that gamma-toxin exhibits specific endonuclease activity for the mcm5s2U-containing tRNA-Glu(UUC) across a spectrum of eukaryotic organisms. Moreover, these studies provide the first experimental evidence that the mcm5s2U modification is conserved in the tRNA of single-celled protists as well as non-mammalian vertebrates such as frogs.

### Gamma-toxin is specific for tRNAs containing mcm5s2U but not similar derivatives

In addition to tRNA-Glu-UUC, the wobble position of eukaryotic tRNA-Arg-UCU, Gln-UUG and Lys-UUU are also known to contain mcm5s2U (Kuntzel et al. 1975; Keith 1984; Chen et al. 2009; Fu et al. 2010a; Songe-Moller et al. 2010; van den Born et al. 2011; Fernandez-Vazquez et al. 2013; Karlsborn et al. 2014a; Yoshida et al. 2015). Gamma-toxin has been shown to cleave these mcm5s2U-containing tRNAs in *S. cerevisiae*, albeit at lower efficiency (Lu et al. 2005) (Supplemental Figure 1). Using RNA isolated from human cells, we find that gamma-toxin can also cleave human tRNA-Arg-UCU, Gln-UUG and Lys-UUU in the anticodon loop (Figure 2, mcm5s2U). These results indicate that gamma-toxin can be used to monitor or verify the presence of mcm5s2U in multiple eukaryotic tRNAs.

**Figure 2.**
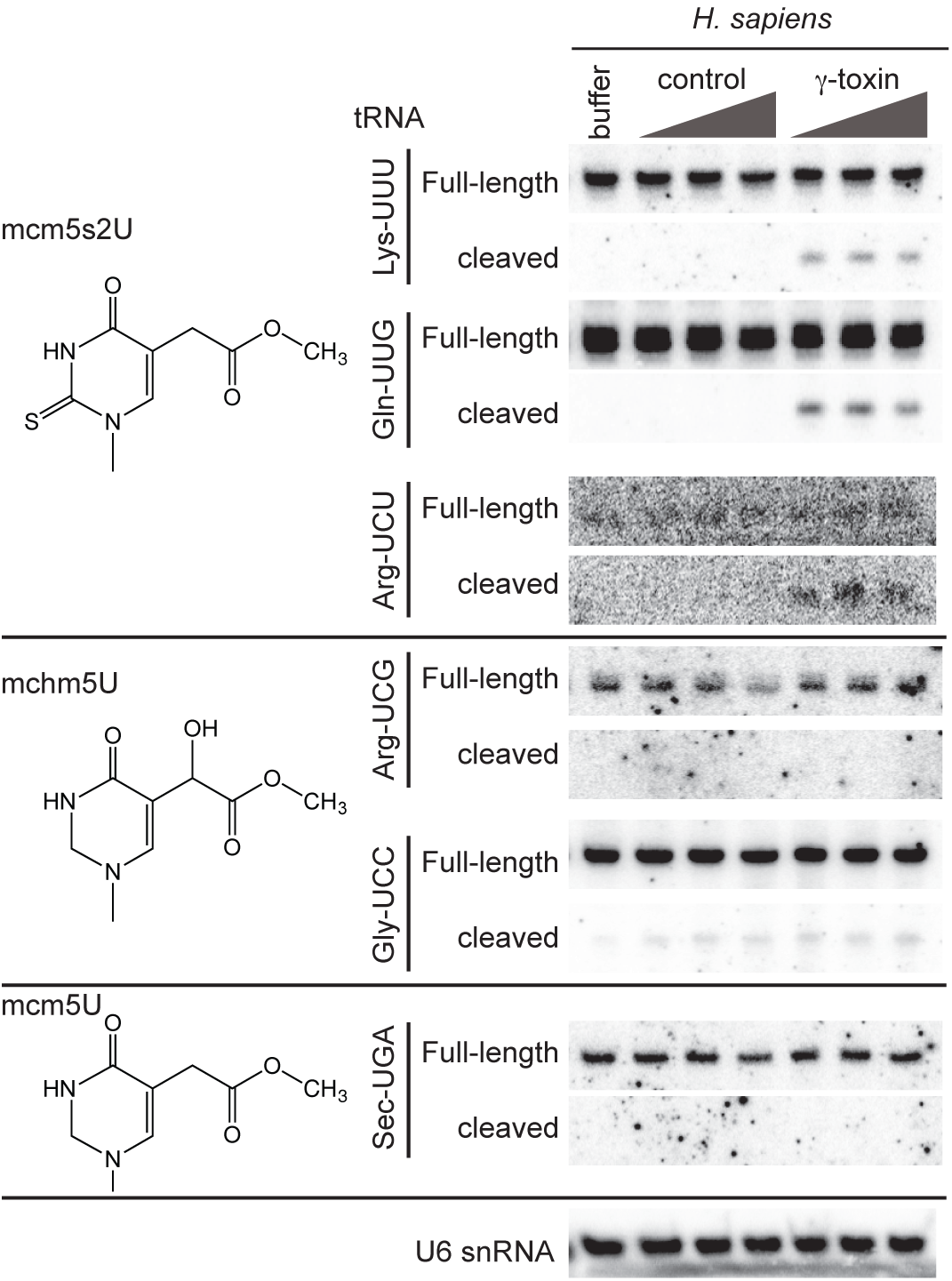
Gamma-toxin is highly specific for tRNAs containing the mcm5s2U modification. TotalRNA extracted from human HEK293T cells was incubated with either buffer, or increasing amounts of a control purification or gamma-toxin followed by Northern blot analysis with probes against the indicated tRNAs. The modification at the Wobble uridine of each tRNA is denoted on the left. U6 snRNA serves as a loading and specificity control.

Besides mcm5s2U-containing tRNAs, we also tested whether gamma-toxin could cleave human tRNAs known to contain modified wobble uridines that differ slightly in chemical composition from mcm5s2U. Previous studies have shown that the mcm5U side chain of mammalian tRNA-Arg-UCG and tRNA-Gly-UCC undergoes enzymatic oxidation to generate the hydroxylated modification, 5-methoxycarbonylhydroxymethyluridine (mchm5U) (Fu et al. 2010b; van den Born et al. 2011). However, gamma-toxin did not exhibit any detectable cleavage activity on either tRNA-Arg-UCG or Gly-UCC (Figure 2, mchm5U). Gamma-toxin was also inactive for cleaving tRNA-Sec-UGA, which contains the 5-methoxycarbonylmethyl-2′-O-methyluridine (mcm5Um) modification at the wobble uridine position (Figure 2, mcm5U). These results demonstrate that gamma-toxin is highly specific for the mcm5s2U modification with lack of the thiol or addition of a hydroxyl group abolishing recognition and cleavage.

### Using gamma-toxin cleavage to test the role of eukaryotic proteins in mcm5s2U formation

In *S. cerevisiae*, the heterodimeric Trm9-Trm112 complex catalyzes the final formation of the methoxycarbonylmethyl side chain in mcm5s2U (Mazauric et al. 2010; Chen et al. 2011a). Trm9 is the catalytic methyltransferase subunit while Trm112 is required for enzymatic activity by serving as a platform for tRNA interaction and cofactor binding (Letoquart et al. 2015; Bourgeois et al. 2017). Numerous homologs of *S. cerevisiae* Trm9 and Trm112 have been identified in eukaryotic and archaeal species (Grosjean et al. 2010; Leihne et al. 2011; Phillips and de Crecy-Lagard 2011; Pastore et al. 2012; Towns and Begley 2012; Zdzalik et al. 2014). However, the activities of the predicted enzymes remain unverified in most cases. Thus, we investigated whether the gamma-toxin assay could be used to validate the methyltransferase activities of Trm9 and Trm112 homologs. For this assay, we used total RNA extracted from *S. cerevisiae trm9*Δ cells as a substrate since it contains the available cm5s2U modification in tRNA as the final methyl acceptor (Mazauric et al. 2010; Chen et al. 2011a). The *trm9*Δ cellular RNA is first incubated with a putative tRNA methyltransferase in the presence of *S*-adenosyl-methionine followed by incubation with gamma-toxin. The level of gamma-toxin cleavage of tRNA is then used as an indicator of mcm5s2U formation catalyzed by the Trm9-Trm112 complex (Figure 3A).

**Figure 3.**
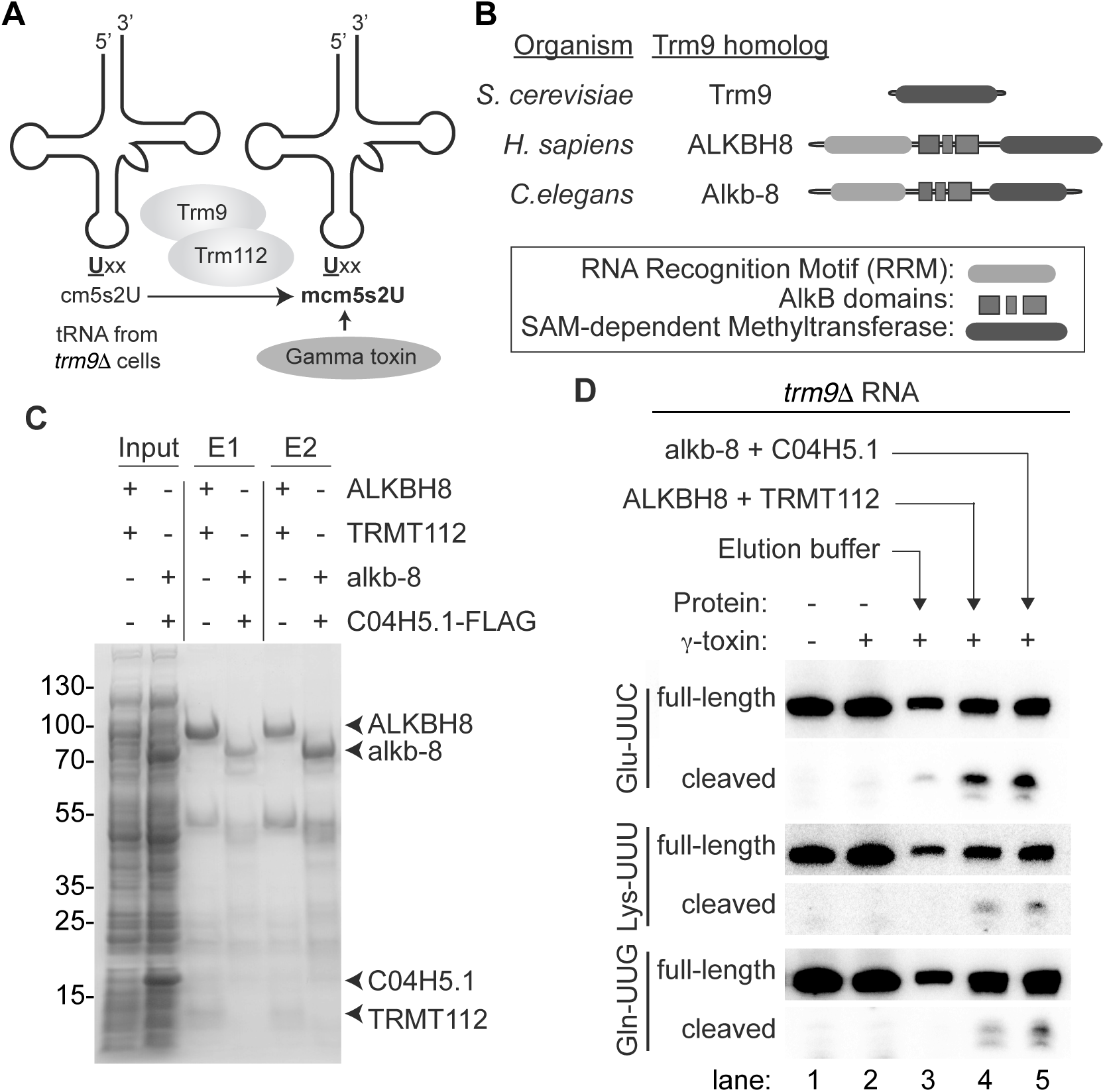
Using the gamma-toxin assay to validate the biochemical activity of putative Trm9-Trm112 homologs. (A) Schematic of gamma-toxin assay to characterize Trm9-Trm112 enzymes for methyltransferase activity. The tRNA of *S. cerevisiae trm9Δ* strains lack the mcm5s2U modification and instead harbours the cm^5^U modification that can be methylated *in vitro* by purified Trm9-Trm112 enzyme complexes. Formation of mcm5s2U can be subsequently detected by the Gamma-toxin assay. (B) Schematic of *S. cerevisiae* Trm9p along with the *H. sapiens* and *C. elegans* homologs, ALKBH8 and Alkb-8, respectively. Both human and *C. elegans* homologs contain a SAM-dependent methyltransferase domain in addition to a RNA recognition motif (RRM) and AlkB dioxygenase domain. (C) Purified protein expressed in *E. coli* cells along with cellular inputs were fractionated by SDS-PAGE and visualized by Coomassie stain. Arrows denote respective proteins. (D) Total RNA extracted from the yeast *trm9Δ* line was incubated with either elution buffer, human ALKBH8-TRMT112 or *C. elegans* alkb-8-C04H5.1 and then subjected to the gamma-toxin assay.

To explore the use of the gamma-toxin assay for identifying eukaryotic proteins involved in mcm5s2U formation, we expressed and purified putative Trm9 and Trm112 homologs encoded by the *C. elegans* genome. Sequence alignment has revealed potential *C. elegans* homologs of yeast Trm9 and Trm112 that are encoded by the Alkb-8 and C04H5.1 genes, respectively (Vilella et al. 2009; Pastore et al. 2012). The Alkb-8 gene product encodes a 591-amino acid protein that displays 32% identity and 47% similarity to the methyltransferase domain of *S. cerevisiae* Trm9 (Figure 3B). In addition to the SAM-dependent methyltransferase domain, *C. elegans* Alkb-8 contains an AlkB dioxygenase domain and RNA recognition motif (Pastore et al. 2012). The *C. elegans* C04H5.1 gene contains an open reading frame encoding a polypeptide of 125 amino acid residues with 23% identity and 55% similarity to *S. cerevisiae* Trm112. However, neither the gene product of Alkb-8 nor C04H5.1 have been characterized in terms of biochemical activity so their status as functional Trm9 and Trm112 homologs remain to be shown.

We co-expressed the *C. elegans* homologs of Trm9 and Trm112 in bacteria followed by metal affinity purification. Alkb-8 was tagged with a hexa-histidine tag for purification purposes while the *C. elegans* Trm112 homolog was epitope-tagged with a 3xFLAG tag for detection. As a positive control, we compared the *C. elegans* proteins to purified human ALKBH8-TRMT112, which forms a heterodimeric complex that has previously been shown to exhibit methyltransferase activity to form mcm5s2U (Fu et al. 2010a). As expected, eluted fractions of affinity purified human ALKBH8 led to the co-purification of TRMT112 (Figure 3C, E1 and E2, +ALKBH8+TRMT112, Supplemental Figure 2). We also detected the co-precipitation of the *C.elegans* C04H5.1 protein with Alkb-8, thereby demonstrating the formation of a heterodimeric complex between the predicted C. elegans Trm9 and Trm112 homologs (Figure 3C, Supplemental Figure 2).

The purified *C. elegans* Trm9-Trm112 complexes were then tested for methyltransferase activity on tRNA isolated from Trm9Δ yeast strains. As noted previously, we detected only slight cleavage of either tRNA-Glu-UUC, Lys-UUU or Gln-UUG when pre-incubated with gamma- toxin due to the lack of mcm5s2U in *trm9*Δ tRNA that is critical for recognition by gamma-toxin (Figure 3D, lanes 1 and 2). In contrast, we could detect gamma-toxin cleavage of tRNA-Glu-UUC, Lys-UUU or Gln-UUG if the *trm9*Δ RNA was first pre-incubated with human ALKBH8-TRMT112 (Figure 3D, ALKBH8+TRMT112, lane 4). This result indicates that recombinant human ALKBH8-TRMT112 enzyme has generated the final mcm5s2U modification in the tRNA isolated from *trm9*Δ cells that is now a substrate for gamma-toxin cleavage *in vitro*. Notably, we could detect robust cleavage of tRNA-Glu-UUC, Lys-UUU or Gln-UUG by gamma-toxin after pre-incubation with the reconstituted *C. elegans* Trm9-Trm112 complex (Figure 3D, alkb-8+C04H5.1, lane 5). Thus, the gamma-toxin assay provides the first demonstration that *C. elegans* Alkb-8 and C04H5.1 (Trm112) form an enzymatically-active tRNA methyltransferase complex that can catalyze the formation of mcm5s2U. Altogether, these results highlight the use of gamma-toxin as a tool to discover and probe the methyltransferase activity of uncharacterized Trm9 and Trm112 homologs from diverse eukaryotic organisms.

### Gamma toxin-coupled real-time qRT-PCR to quantify mcm5s2U levels

Thus far, we have detected gamma-toxin cleavage using Northern blotting with radiolabeled probes. While Northern blotting provides a sensitive method for RNA detection, the protocol requires expertise in RNA electrophoresis, blotting equipment and radioactive probe labeling. Thus, we developed a quantitative real-time qRT-PCR approach to measure gamma-toxin cleavage of tRNA as a rapid, non-radioactive alternative to Northern blot analysis for monitoring the levels of mcm5s2U. For this method, we treated sample RNAs with gamma-toxin followed by reverse transcription using a primer downstream of the tRNA anticodon sequence to generate a mixture of cDNA products representing full-length or truncated cDNA products (Figure 4A). Next, we PCR amplified the mixture of cDNA products using primers spanning the gamma-toxin cleavage site. Since the presence of mcm5s2U would lead to gamma-toxin cleavage of tRNA, the expectation would be to detect less cDNA product by quantitative PCR. Conversely, a decrease in mcm5s2U modification would lead to less gamma-toxin cleavage and more full-length cDNA product detectable by PCR. We then normalized the signal to rRNA to control for the amount of RNA within the same sample.

**Figure 4.**
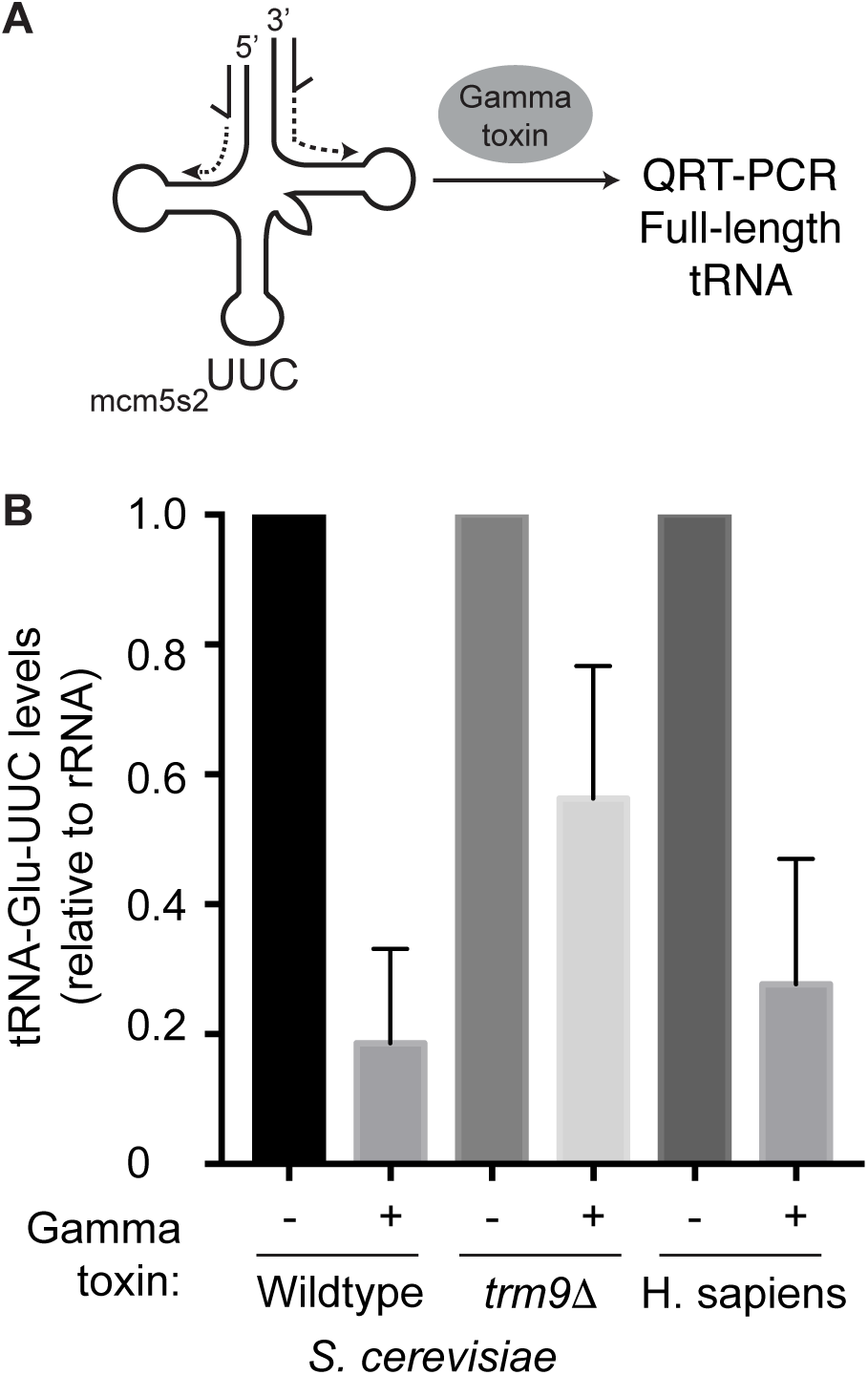
Quantitative detection of gamma-toxin cleavage of mcm5s2u-containing tRNAs using qRT-PCR. (A) Schematic of qRT-PCR assay for detecting tRNA cleavage by gamma-toxin. The location of PCR primers for amplification of full-length tRNA-Glu-UUC is denoted by half-arrows. (B) Real-time PCR analysis of cDNA generated from reverse transcription of total RNA isolated from the indicated yeast strain or human cells that were either untreated or pre-treated with gamma-toxin. The total amount of RNA was normalized using 25S rRNA (yeast) or 5.8S (human) and expressed relative to the untreated total RNA sample of each organism. All assays were performed in triplicate on multiple independent samples and repeated greater than 3x to ensure reproducibility.

As a benchmark comparison to Northern blotting, we tested the detection sensitivity of qRT-PCR for monitoring mcm5s2U in *S. cerevisiae* tRNA after gamma-toxin cleavage. We found that qRT-PCR could detect an ~80% decrease in full-length tRNA-Glu-UUC after treatment with gamma-toxin, comparable to the detection sensitivity by Northern blot analysis (Figure 4B). We also used qRT-PCR to monitor gamma-toxin cleavage of tRNA-Glu-UUC isolated from *trm9*Δ strains, which are known to lack mcm5s2U and cannot be cleaved by gamma-toxin. Indeed, we found that the levels of full-length tRNA-Glu-UUC after gamma-toxin treatment was higher for *trm9*Δ strains versus WT yeast cells (Figure 4B, compare WT versus *trm9*Δ + gamma-toxin). To demonstrate the utility of the assay beyond *S. cerevisiae*, we also monitored mcm5s2U levels in human tRNA-Glu-UUC with our integrated gamma-toxin qRT-PCR approach. We found that qRT-PCR could detect a ~70% decrease in tRNA-Glu-UUC levels after gamma-toxin treatment (Figure 4B). Thus, the combination of gamma-toxin cleavage with qRT-PCR provides a rapid and sensitive method to probe the levels of mcm5s2U levels in different genetic backgrounds or conditions.

### Future Applications

Here, we have shown that recombinant gamma-toxin displays robust and specific endonuclease activity to cleave mcm5s2U-containing tRNAs in diverse eukaryotic organisms. These results demonstrate that gamma-toxin can be applied as a tool for investigating the levels of mcm5s2U modification in a variety of model eukaryotic systems including *D. melanogaster*, *C.elegans* and mammalian cells. The use of gamma-toxin technique is straightforward by using short incubation, Northern blotting and detection of cleaved products. The gamma-toxin protocol can also be combined with real-time qRT-PCR, thus providing a rapid and sensitive alternative to Northern blotting for quantifying the presence of mcm5s2U in the tRNA of eukaryotes. Thus, gamma-toxin represents an efficient and specific detection method for assaying the mcm5s2U tRNA modification in all known tRNAs containing the mcm5s2U modification.

In addition to monitoring the levels of mcm5s2U in tRNA, the gamma-toxin assay can be used to identify and validate putative eukaryotic proteins that play a role in mcm5s2U formation. Indeed, we have used the gamma assay to demonstrate the methyltransferase activity of human and worm homologs of Trm9-Trm112. We envision that the gamma-toxin assay can be extended to discover putative homologs of the Elongator and thiolase complexes as well as novel subunits involved in mcm5s2U formation. Moreover, the gamma-toxin assay can be used as an assay to distinguish Trm9 paralogs with different targets such as the case of ALKBH8 and KIAA1457 in humans (Fu et al. 2010a; Songe-Moller et al. 2010; Begley et al. 2013). Furthermore, the gamma-toxin assay could be applied to the numerous predicted Archaeal homologs of Elongator and Trm9 that have not been characterized in terms of their final biochemical activities (Grosjean et al. 2008; Phillips and de Crecy-Lagard 2011; Naor et al. 2012).

Finally, our findings suggest that gamma-toxin could be expressed in heterologous organisms as a way to test translation dependent upon tRNAs containing the mcm5s2U modification. For example, gamma-toxin could be transiently expressed in mouse or human cells to cleave endogenous mcm5s2U-containing tRNAs followed by proteomic analysis as a tunable method to analyze the role of mcm5s2U modifications in translation. Altogether, these studies will pave the way for understanding the biological role of mcm5s2U modification and their ubiquitous nature in eukaryotes.

## MATERIAL AND METHODS

### Protein expression constructs

The protein expression construct for gamma-toxin was provided by the Shuman Lab (pET28-His10Smt3-gamma-toxin-C13A-C177A-C231A) (Keppetipola et al. 2009). The open reading for human TRMT112 was PCR amplified from cDNA plasmid HsCD00323319 (PlasmID Repository, Harvard Medical School) and cloned into the BglII-KpnI sites of pET-Duet1 (EMD Millipore) to generate pET-Duet1-TRMT112. The open reading frame for human ALKBH8 was PCR amplified from pcDNA3.1-FLAG-ALKBH8 (Fu et al. 2010a) and cloned into pET-Duet-TRMT112 using SacI-SalI. The dual ALKBH8-TRMT112 construct was generated by cloning NotI-XhoI fragments from pDuet-TRM112 into the identical sites of pET-Duet1-ALKBH8. The open reading frames of *C. elegans* Alkb-8 and TRM-112 were PCR amplified from the cDNA clones, C14B1.10 and C04H5.1, respectively (PlasmID Repository, Harvard Medical School). The Alkb-8 and TRM-112 inserts were cloned into pET-Duet1 using BglII-KpnI and SacI-HindIII, respectively. All inserts were sequenced and verified for the absence of mutations.

### Protein expression and purification

The pET28a bacterial expression vector containing either N-terminal His-tagged gamma-toxin or control vector was transformed into BL21 (DE3)-RIPL *E.coli* cells (Agilent Technologies) and grown in 50 ml of Luria-Bertani media containing kanamycin and chloramphenicol to an OD_600_ of 0.6 - 0.8. Protein expression was induced by the addition of Isopropyl β-D-1-thiogalactopyranoside (IPTG) for 18 hours at 16°C at a final concentration of 0.4 M. Cells were then pelleted at 4000 x g for 10 minutes and resuspended in 4 ml of Bacterial Lysis Buffer (20 mM Tris, 5% glycerol, 0.3%Triton, 1 mM DTT, 0.1 mM PMSF, 150 mM NaCl, 25 mM imidazole). Cells were lysed via sonication and cellular debris was pelleted at 20,000 x g for 30 mins at 4°C. Cellular extracts were incubated with HisPur Ni-NTA Resin (Thermo Fisher Scientific) and rotated at 4°C for two hours before being washed 3 times in the aforementioned lysis buffer. Protein was eluted using a buffer containing 300 mM imidazole, 200 mM NaCl, 20 mM Tris, 5% glycerol, 0.3% Triton, and PMSF. Purified protein was visualized on a 15% SDS-PAGE gel stained with Coomassie protein stain. BL21 (DE3)-RIPL *E.coli* cells harboring the pETDuet1 bacterial expression vectors containing either *Homo sapiens* or *Caenorhabditis elegans* Trm9-Trm112 homologs were grown as mentioned above. Protein expression of Human ALKBH8-TRMT112 by IPTG was induced for 15 hours at 20°C. *C. elegans* Alkb-8 and TRM-112 bacterial expression vector was induced for 15 hours at 16°C. Cells were then pelleted at 4000 x g for 10 minutes and resuspended in 4 ml of Bacterial Lysis Buffer (20 mM Tris, 5% glycerol, 0.3%Triton, 1 mM DTT, 0.1 mM PMSF, 150 mM NaCl (200 mM for Human homologs), and 25 mM imidazole). Cells were lysed and proteins were purified as described above for Gamma-toxin.

### RNA isolation

Wildtype and *trm9Δ S. cerevisiae* strains were obtained from the Sia Lab (University of Rochester). Colony PCR was performed to ensure the correct genotype. *S. cerevisiae* strains were grown until OD_600_ of 1.0 and total RNA was purified using the hot acid phenol technique (Collart and Oliviero 2001). *E. coli* RNA was extracted following the RNA*snap* method (Stead et al. 2012). For *Tetrahymena thermophila*, human and mouse samples, RNA was purified directly from cell pellets using TRIzol LS RNA extraction (Invitrogen). Tetrahymena strain SB210 were grown in standard proteose peptone growth media to mid-log phase before harvesting. Human embryonic kidney (HEK) 293T cells were grown in DMEM supplemented with 10% fetal bovine serum (FBS), 1X antibiotics and 1X Glutamax at 37°C with 5% CO_2_. Mouse embryonic fibroblast (MEF) were grown in DMEM supplemented with 15% FBS at 37°C with 5% CO_2_.

Whole organism RNA extraction was performed on *C. elegans* and *D. melanogaster*. Vials containing adult *D. melanogaster* were frozen at -80°C for 10 minutes. 25 mg of frozen adult flies were homogenized with a plastic pestle in a 1.5 mL microfuge tube while adding 250 μl of TRIzol reagent at a time until 1 ml was added. RNA was then extracted via an adapted Trizol extraction protocol (Bogart and Andrews 2006). For extraction of *C. elegans* RNA, a confluent 60 mm plate of adult worms was scraped into a 1.5 mL microfuge tube and resuspended in 1 ml of RNAse-free water. Tubes were centrifuged at 14,000 x g for 10 minutes and 750 μl water was removed. The worm pellet was resuspended in the remaining 250 μl water and processed using TRIzol LS.

For *X. laevis* samples, the liver from one adult *X. laevis* was surgically extracted, flash frozen in liquid nitrogen and stored immediately at -80°C. One quarter of an adult liver was used to extract RNA at a given time. While still frozen, one fourth of the liver was cut into multiple slices, transferred to a 1.5 mL microfuge tube and homogenized with 750 μl of TRIzol using a plastic pestle. Following this step, TRIzol purification was followed according to standard protocol.

### Gamma-toxin assay

For each species tested, 5 μg of total RNA was incubated for 10 minutes at 30°C with increasing volumes of purified gamma-toxin or control eluate in 10 mM Tris-HCl (pH 7.5), 10 mM MgCl_2_, 50 mM NaCl and 1 mM dithiothreitol (DTT, pH 7.5). For the yeast and worm blots, the gamma-toxin was diluted by a factor of 12.5. Each sample was run on a 10% polyacrylamide, 7 M urea gel and transferred to an Amersham Hybond-XL membrane (GE Healthcare). Oligonucleotides used to detect tRNAs are listed in Supplemental Table 1. The oligos were radiolabeled by T4 polynucleotide kinase (NEB) with adenosine [γ^32^P]-triphosphate (6000 Ci/mmol, Amersham Biosciences) following standard protocols. Northern blots were visualized by Phosphor-Imager analysis. Blots were subsequently stripped via two incubations at 80°C for 20 minutes in a buffer containing 0.15 M NaCl, 0.015 M Na-citrate and 0.1% SDS.

### Reconstitution and activity assay of Trm9-Trm112 homologs

Purified human and *C. elegans* Trm9-Trm112 homologs were mixed with 5 μg total RNA harvested from *trm9Δ* yeast deletion strain along with RNAse inhibitor, 160 μM *S-*adenosylmethionine, 250 mM Tris (pH 7.5), 25 mM CaCl_2_, 400 μM NH_4_Ac, 500 μM MgCl_2_ and 100 μM EDTA. In place of purified protein, water was used as a control to ensure non-specific cleavage was not occurring throughout the process. Reactions were incubated at 37°C for two hours before being RNA purified through Zymo-Spin RNA Clean and Concentrator IC columns (Zymo Research). The gamma-toxin assay was then performed on each purified RNA sample as described above.

### QRT-PCR assay

5 μg of total RNA underwent gamma-toxin treatment as previously described. RNA was extracted via the TRIzol method and resuspended in 15 μl of RNase-free water. 1 μg of purified RNA was used to create cDNA using SuperScript IV reverse transcriptase (ThermoFisher Scientific). cDNA was created for both tRNA-Glu for both yeast and human as well as an rRNA normalizing control (25S for yeast and 5.8S for human). RNA was first incubated with 10 mM dNTPs and 10 μM reverse primer (Listed in Supplemental Table 1). Reactions were incubated at 65°C for 5 mins and then left on ice for 2 minutes before the addition of RNAse Inhibitor (Thermo Fisher Scientific), 100 mM DTT, SuperScript IV reverse transcriptase and its accompanying buffer at 50°C for an hour followed by 80°C heat inactivation for 10 minutes. Each reaction was then cleaned up using the Qiagen PCR clean-up kit. 2 μl of cDNA was added to a reaction with SYBR Green Real-time PCR Master Mixes (Thermo Fisher) and 10 μM of forward and reverse primers (Supplemental Table 1). Quantitative PCR analysis was conducted with a Biorad PCR thermocycler. Each reaction was performed in triplicate. Relative tRNA levels were calculated using the ΔΔCt method and normalized to either yeast 25S rRNA or human 5.8S rRNA.

## SUPPLEMENTAL MATERIAL

Supplemental Table 1. Oligonucleotides used in this study.

Supplemental Figure 1. Gamma-toxin cleavage of *S. cerevisiae* tRNA-Gln-UUG and Lys-UUU.

Supplemental Figure 2. Immunoblot verification of Trm9-Trm112 copurification.

## ACKNOWLEDGEMENTS

We thank Stewart Shuman (Memorial Sloan Kettering) for the gamma-toxin expression plasmid, Elaine Sia and Christopher Prevost (University of Rochester) for yeast strains, Jacques Robert (University of Rochester Medical Center) for *X. laevis* liver samples, Douglas Portman (University of Rochester Medical Center) for *C. elegans*, Danna Eickbush and Amanda Larracuente (University of Rochester) for *D. melanogaster*, Suzanne Lee (Washington State University) for *Tetrahymena thermophila* cells and Yan Li for cloning of *C. elegans* Trm9-Trm112.

---

**Supplemental Table 1.**
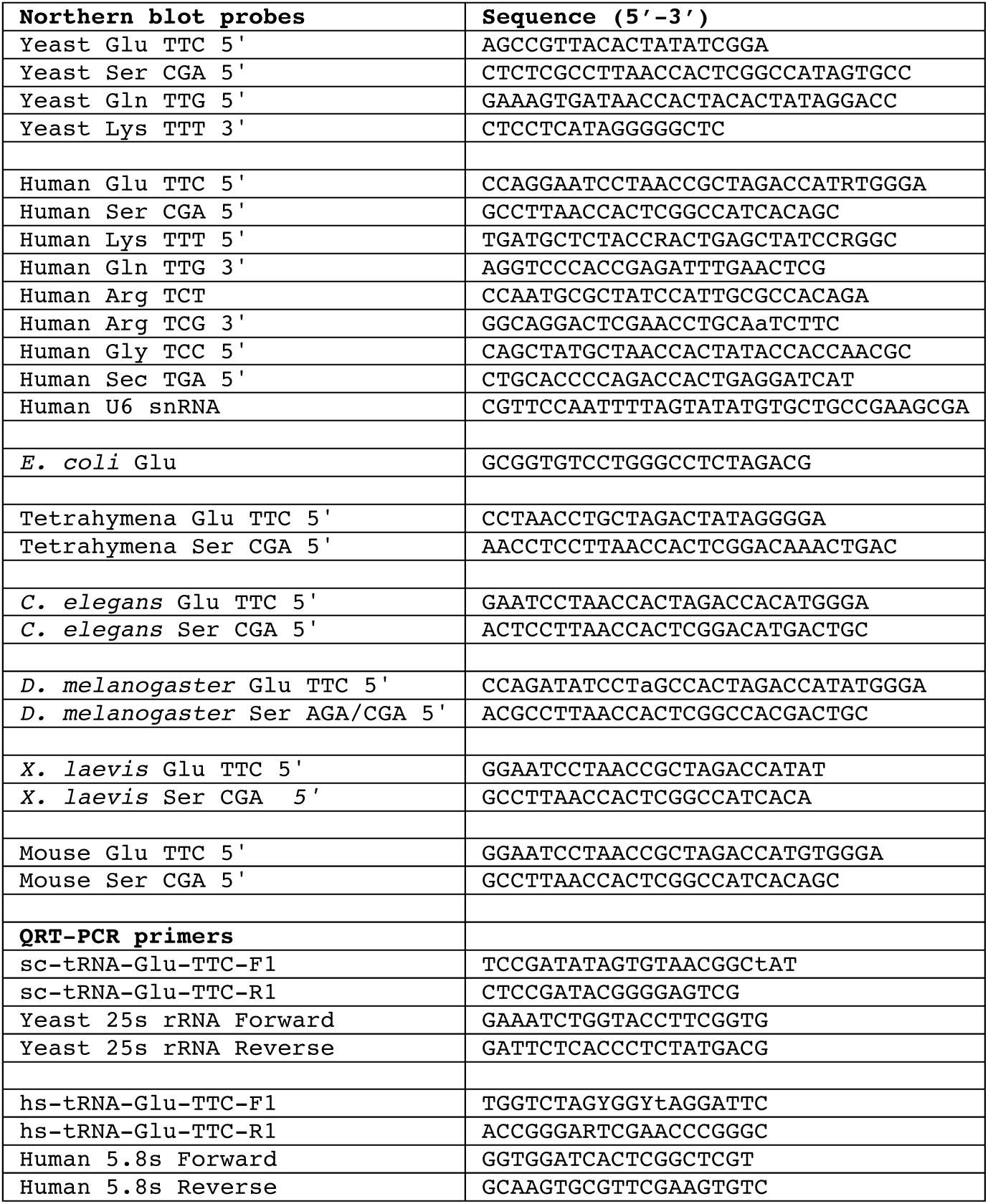
Oligonucleotides

**Supplemental Figure 1.**
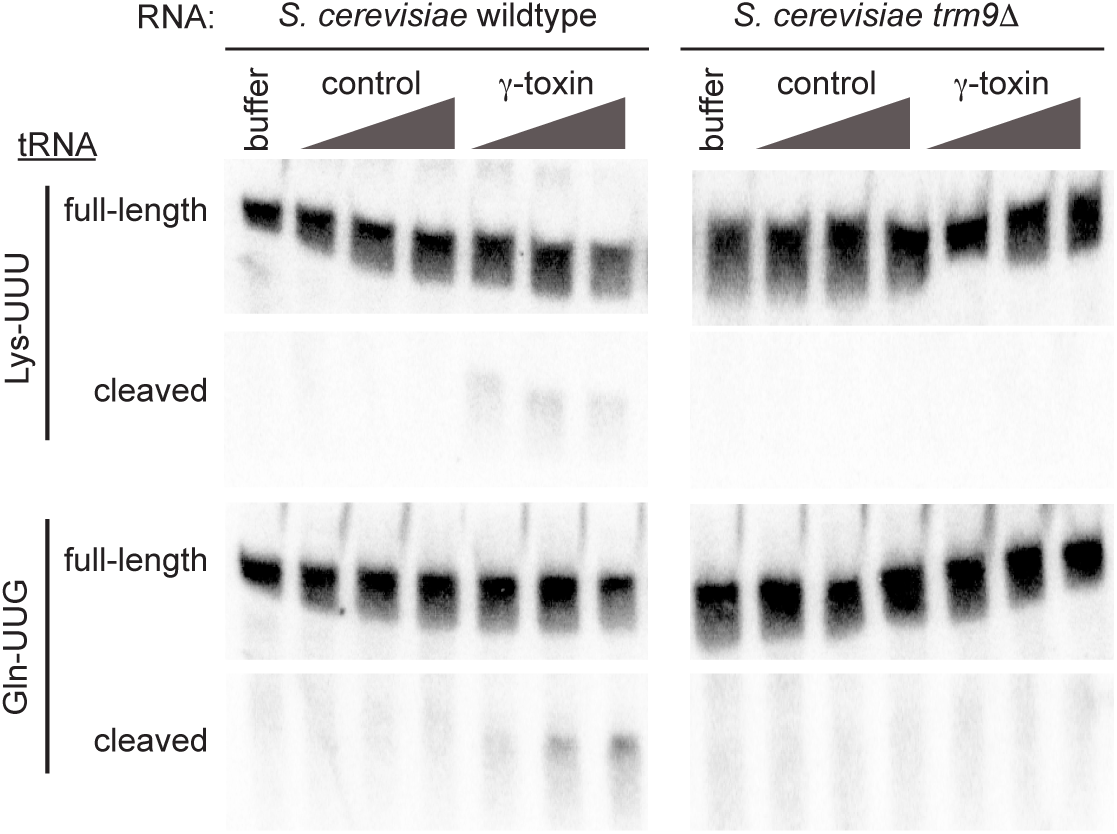
Gamma-toxin cleavage of *S.cerevisiae* tRNA-Gln-UUG and Lys-UUU. Northern blots of *S. cerevisiae* Wild-type and trm9delta RNA incubated with buffer alone or increasing amounts of control purification or purified gamma-toxin. The blot was hybridized with probes against tRNA-Lys-UUU and tRNA-Gln-UUG.

**Supplemental Figure 2.**
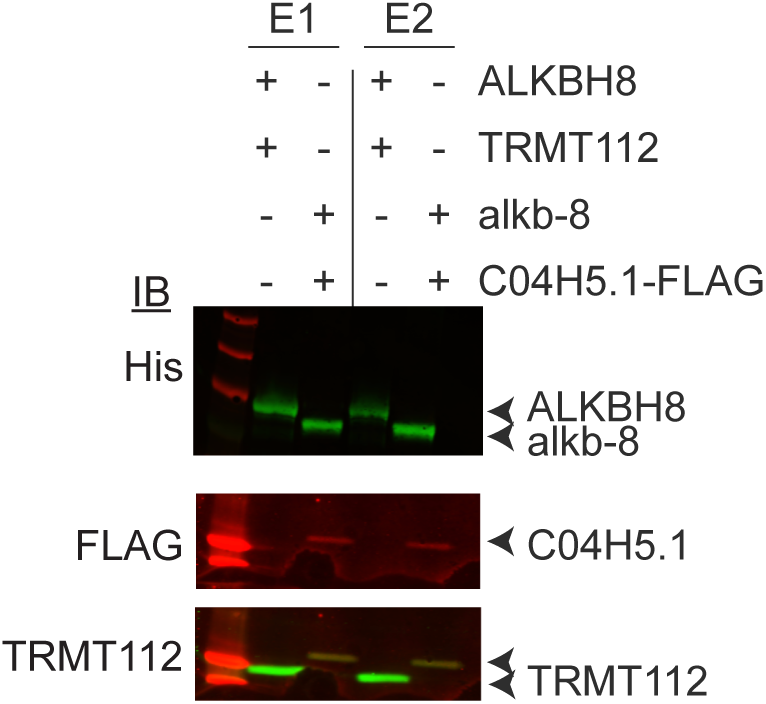
Immunoblot verification of Trm9-Trm112 copurification. Elutions 1 and 2 of purified protein expressed in *E. coli* bacterial cells were visualized by western blotting and antibodies against hexa-histidine tag, 3X-FLAG tag, and TRMT112 were used.

